# Generation of nanobodies acting as silent and positive allosteric modulators of the α7 nicotinic acetylcholine receptor

**DOI:** 10.1101/2023.01.03.522562

**Authors:** Qimeng Li, Ákos Nemecz, Gabriel Aymé, Gabrielle Dejean de la Bâtie, Marie S Prevost, Stéphanie Pons, Nathalie Barilone, Rayen Baachaoui, Uwe Maskos, Pierre Lafaye, Pierre-Jean Corringer

## Abstract

The α7 nicotinic acetylcholine receptor (nAChR), a potential drug target for treating cognitive disorders, mediates communication between neuronal and non-neuronal cells. Although many competitive antagonists, agonists, and partial-agonists have been found and synthesized, they have not led to effective therapeutic treatments. In this context, small molecules acting as positive allosteric modulators binding outside the orthosteric, acetylcholine, site have attracted considerable interest. Two single-domain antibody fragments, C4 and E3, against the extracellular domain of the human α7-nAChR were generated through alpaca immunization with cells expressing a human α7-nAChR/mouse 5-HT_3_A chimera, and are herein described. They bind to the α7-nAChR but not to the other major nAChR subtypes, α4β2 and α3β4. E3 acts as a slowly associating positive allosteric modulator, strongly potentiating the acetylcholine-elicited currents, while not precluding the desensitization of the receptor. An E3-E3 bivalent construct shows similar potentiating properties but displays very slow dissociation kinetics conferring quasi-irreversible properties. Whereas, C4 does not alter the receptor function, but fully inhibits the E3-evoked potentiation, showing it is a silent allosteric modulator competing with E3 binding. Both nanobodies do not compete with α-bungarotoxin, localizing at an allosteric extracellular binding site away from the orthosteric site. The functional differences of each nanobody, as well as the alteration of functional properties through nanobody modifications indicate the importance of this extracellular site. The nanobodies will be useful for pharmacological and structural investigations; moreover, they, along with the extracellular site, have a direct potential for clinical applications.

## Introduction

Nicotinic acetylcholine receptors (nAChRs) belong to the pentameric ligand-gated ion channel family and play a key role in neuronal communication as well as in non-neuronal cells such as immune and epithelial cells. The major nAChRs in the brain and at the periphery are the homomeric α7-nAChR, and the heteromeric α4β2- and α3β4-nAChRs, a fraction of these later incorporating the accessory α5 or β3 subunits [1]. Acetylcholine (ACh) binding promotes a global reorganization in nAChRs, whereupon their intrinsic channel opens, while the prolonged binding of ACh promotes a second reorganization, where the channel closes in what is termed the desensitized state. Among nAChRs, α7-nAChR displays unique properties, including a low probability of channel opening and rapid desensitization [2].

The α7-nAChR has attracted considerable interest and been pursued as a potential therapeutic target for numerous indications [3]. The α7-nAChR is abundant in brain regions such as the hippocampus and the prefrontal cortex that are important for cognitive functions. Therefore, drugs that activate or potentiate the receptor have been shown to be effective in preclinical models for cognitive disorders [4]. Additionally, several therapeutics were tested through clinical trials in the context of Alzheimer’s and Parkinson’s diseases, as well as schizophrenia [5]. However, as of yet there has not been any approval for clinical use, either due to lack of efficacy or to adverse effects [6]. The α7-nAChR is also an essential component of the cholinergic anti-inflammatory pathway, specifically its activation through excitation of the vagus nerve triggers release of anti-inflammatory cytokines [7]. Of note, the α7-nAChR is not only found as homopentamers in the brain, but also as heteropentamers in complex with the β2 subunit [8], as well as with the dupα7 subunit, which is a truncated subunit, associated with neurological disorders, lacking part of the N-terminal extracellular ligand-binding domain [9].

A lot of effort has been dedicated to developing small molecules specifically targeting the α7-nAChRs. Each nAChR subunit within the pentamer is composed of an extracellular domain (ECD) folded as a β sandwich, a transmembrane domain (TMD) consisting of four α-helices, and an intracellular domain (ICD) consisting of two helices and a variably sized poorly resolved domain connecting the two [10, 11]. The endogenous neurotransmitter’s (ACh) binding sites, also called orthosteric sites, are located at all of the subunit interfaces within the ECD of the homomeric α7-nAChR. Agonists, partial agonists, and antagonists all bind at the orthosteric site and were the first therapeutic focus, whereas negative (NAM) and positive (PAM) allosteric modulators binding outside of this site have also actively been investigated more recently [3]. Indeed, the very rapid desensitization of α7-nAChRs is expected to strongly limit the efficacy of conventional agonists, while allosteric modulators can potentially overcome this issue. In addition, PAMs and NAMs are expected to better maintain the spatio-temporal characteristics of endogenous ACh activation and to target non-conserved sites, increasing the chemical diversity of active compounds.

In chronological order, calcium was first identified as a PAM [12, 13] binding in the lower part of the ECD [10, 14]. Ivermectin was then identified as a PAM binding in the TMD [15]. Ivermectin can be classified as a type I PAM, potentiating the ACh-elicited current at the peak of the electrophysiological response but not impairing the downstream desensitization process. Subsequently, a large series of small molecules binding at the TMD were found to strongly modulate the receptor, as exemplified by the type II PAM: PNU-120596, that not only potentiates the ACh-elicited currents but also inhibits desensitization to a large extent [11, 16]. Additionally, the PAM: 4BP-TQS can even activate the receptor by itself, thereby having both agonistic and modulatory properties outside of the orthosteric site (Ago-PAM) [17]. Finally, several modulatory sites for small fragments were identified at different levels of the ECD but have not, as of yet, been exploited for drug-design purpose [18, 19].

In addition to small molecules, an interest has recently grown around single-domain antibody fragments of camelids, generally termed nanobodies, in developing biotechnologies [20]. Nanobodies correspond to the variable domain (VHH) of the heavy chain-only antibodies expressed in these animals. Moreover, they usually bind to surface cavities [21] and motifs that often reorganize during conformational transitions of the receptor, thereby acting as conformation-specific ligands. In addition, nanobodies have a number of advantages over small molecules, notably: they usually have a high affinity, typically in the nanomolar range, as well as a high specificity conferred by the large surface of the nanobody-antigen interaction. As an example: within the pentameric ligand-gated ion channel family, nanobodies acting as PAMs and NAMs were reported on the serotonin type 3 (5-HT_3_) receptor [22], the GABA_A_ receptor [23], and the bacterial ELIC [24]. The generation and functional characterization of two nanobodies specifically targeting the α7-nAChR is presented herein.

## Results

### 1/Two nanobodies were isolated from alpacas immunized with cells expressing the hα7-nAChR/m5-HT3A chimera

To avoid the time-consuming and expensive production and purification of the α7-nAChR, alpacas were immunized with HEK 293 cells transiently transfected with a cDNA directing the expression of an α7-nAChR/5-HT_3_A chimera. This chimera, where the ECD of the human α7-nAChR is fused to the TMD of the mouse 5-HT_3_A receptor (hα7-nAChR/m5-HT_3_A), is expressed at the surface of the cells at much higher levels than α7-nAChR [25, 26]. In addition, injecting transfected cells ensures that the receptor is correctly folded and inserted in the membrane as compared to a detergent solubilized receptor, thereby increasing the chances of isolating conformation-specific nanobodies. A similar approach has been successful for other membrane proteins such as the metabotropic glutamate receptors [27]. After immunization, the VHH-encoding sequences were amplified from the serum and a phage display library was generated (as described in the methods). In a single round of panning, the library was depleted twice through incubation with non-transfected HEK 293 cells, followed by an incubation with non-transfected Vero cells, after which Vero cells transfected with the hα7-nAChR/m5-HT_3_A chimera were used to select phages binding to the α7-ECD. Isolated clones from the final two rounds of panning were then screened by an ELISA against transfected Vero cells. Seven unique sequences have been obtained from the combination of the second and third rounds of panning.

### 2/Flow-cytometry experiments show that C4 and E3 have robust α7-nAChR binding and immunofluorescence experiments show specificity for the α7-nAChR over α3β4- and α4β2-nAChRs

The clones were produced with a C-terminal Myc and His tag (named VHH in Figure 1b) and tested by flow cytometry, using non-fixed live cells expressing the hα7-nAChR/m5-HT_3_A. They were also tested by immunofluorescence. Among the seven VHHs, C4 and E3 displayed specific labeling both in flow-cytometry and immunofluorescence. Flow cytometry results using the positive control of 0.1mg/mL (∼11.7µM) α-Bungarotoxin (α-Btx) conjugated with AlexaFluor488 resulted in ∼30% of the cells bound and therefore expressing the hα7-nAChR/m5-HT_3_A chimera on the cell surface, whereas 0.02mg/mL (∼1.27µM of each) of the C4 and E3 VHHs, detected using an AlexaFluor488 conjugated anti-C-Myc secondary antibody, bind ∼9% of the cells (Figure 2). Because of the use of a secondary antibody, the VHH samples experienced an extra half hour of washing during the secondary antibody binding compared to the positive, α-Btx, control. VHH F5, representative of the other five VHHs, shows no labeling of either transfected or non-transfected cells.

**Figure 1:**
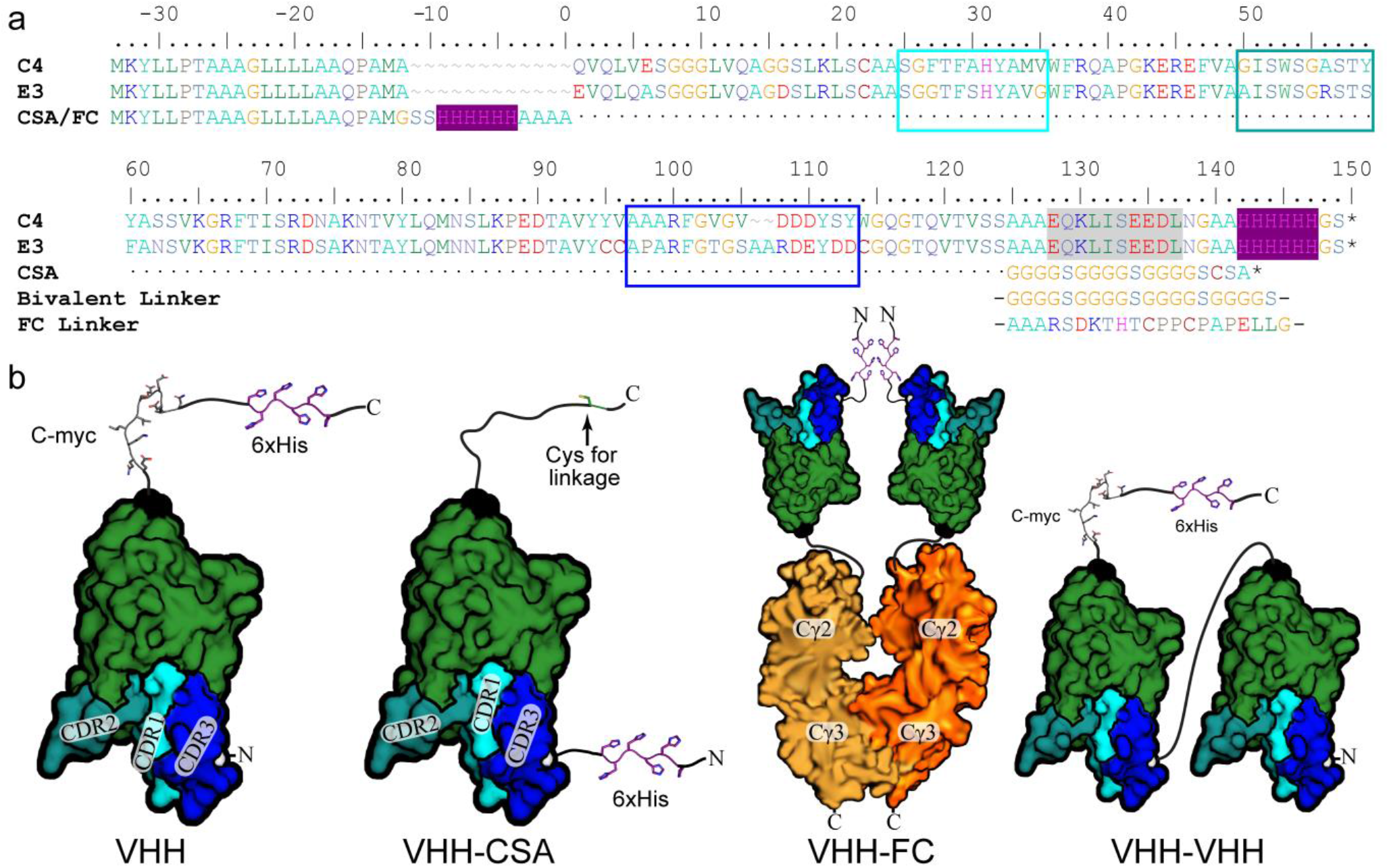
VHH E3 and C4 with Related Constructs. **a** Sequence alignment of E3 and C4 VHHs with numbering starting after the signal peptide, although sometimes incomplete cleavage occurs and the preceding MA of the signal peptide is left attached. The Myc tag is highlighted with gray shading, 6xHis tag highlighted in purple shading, and each CDR is boxed in cyan, teal, and blue (1,2,3 resp.). N- and C-terminal changes for the Fc, CSA, and bivalent constructs are also shown. **b** Structural representation of each variation with the tags labeled and their side-chains represented as sticks, the N- and C-terminals labeled and colored (white and black resp.), as well as the CDRs labeled and color coded (similarly as in **a**).

**Figure 2:**
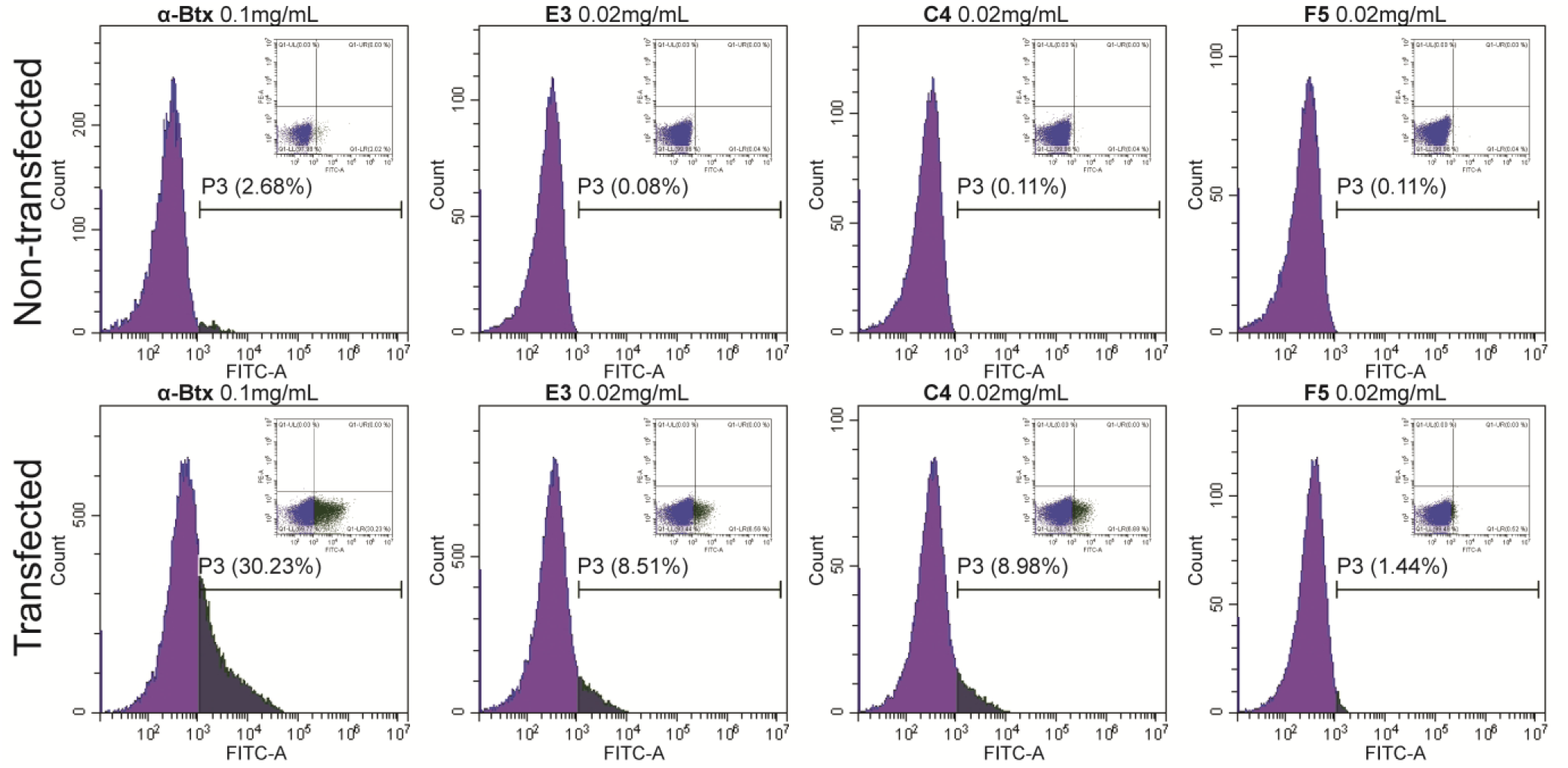
VHH characterization by flow-cytometry. Flow-cytometry histograms with density plots (PE vs. FITC) of the same data inset. P3, the population of cells with bound VHH or α-Btx is labeled. Top row: Non-transfected HEK293 cells labeled with AlexaFluor488 conjugated α-Btx (0.1mg/mL) or the respective VHHs at 0.02mg/mL labeled with a secondary AlexaFluor488 conjugated anti-C-Myc antibody, where F5 is a VHH found not to bind. Bottom row: Corresponding labeling of cells transfected with hα7-nAChR/m5-HT_3_A.

Two other variants of C4 and E3 were generated: 1/VHH-CSA, carrying a C-terminal cysteine, allowing direct fluorescent labeling to avoid the use of secondary antibodies, and 2/VHH-Fc, where VHHs are fused at the C-terminus with a human IgG Fc fragment permitting detection through a highly specific secondary anti-human Ig antibody, as well as inclining the formation of dimeric VHH domains due to the dimeric nature of the IgG-Fc, putatively increasing binding avidity (Figure 1b). The immunofluorescence of both constructs was carried out against non-permeabilized cells expressing the hα7-nAChR (Figure 3). E3-CSA labeled with Alexa488, was also tested using CHO cells stably expressing the hα7-nAChR, and showed specific labeling (Figure 3a). A non-human cell-line, CHO, was chosen for stable hα7-nAChR expression, to avoid cross-reactivity with human proteins in testing of the VHH-CSA construct, whereas a human cell-line, HEK293, was used to test specificity and selectivity of VHH-Fc constructs. The VHH-Fc constructs, combined with an anti-human IgG to label them, displayed a nice decoration of the plasma-membrane for both E3-Fc and C4-Fc on transiently transfected HEK293 cells (Figure 3b), and therefore this construct was used to assess each VHH’s binding specificity. The combination of these results suggests a binding motif in the ECD.

**Figure 3:**
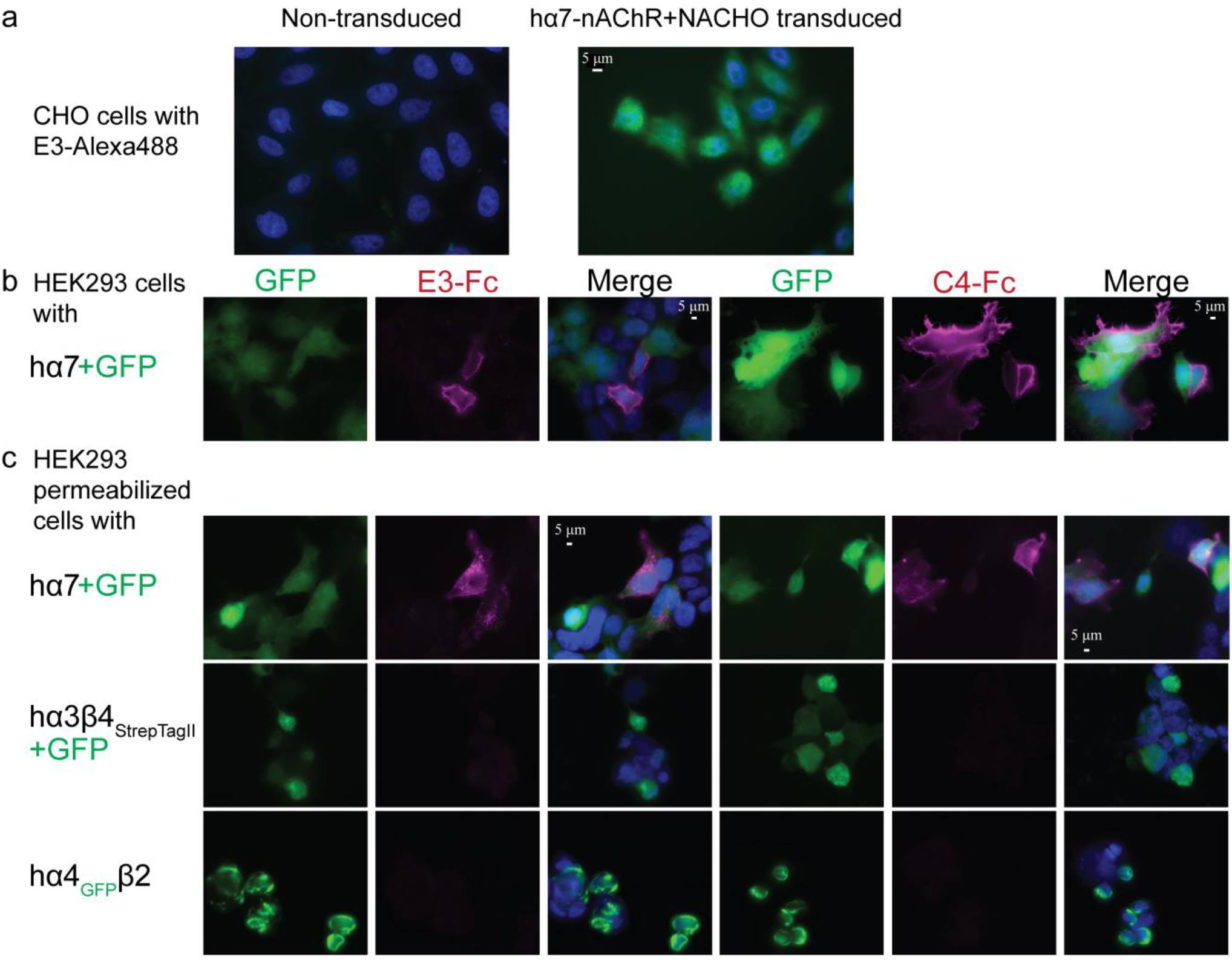
VHH characterization by immunofluorescence. Dapi, shown in blue, stains the cells’ nucleus. Alexa647, a red wavelength, is colored as magenta. Images are representative of at least n=4. **a** Merged images (Dapi+GFP) of CHO cells (left) and with stable expression of hα7-nAChR (right); immunostained with 1µg/mL of conjugated E3-CSA-Alexa488. **b** Images (GFP, Alexa647, Merged [GFP+Alexa647+Dapi]) of non-permeabilized HEK 293 cells transfected with hα7-nAChR immunostained with E3-Fc (left) and C4-Fc (right), demonstrating an extracellular binding. Cytoplasmic GFP indicates efficiently transfected cells. **c** Images (GFP, Alexa647, Merged [GFP+Alexa647+Dapi]) of permeabilized HEK 293 cells expressing hα7-, hα3β4_StrepII_- and hα4_GFP_β2-nAChRs immunostained using E3-Fc (left) and C4-Fc (right). For hα7 and hα3β4_Strep_, cytoplasmic GFP indicates efficiently transfected cells. The nanobodies were detected by an anti-human IgG coupled to Alexa647 (red). Identical exposure times were used to visualize each channel on all conditions.

The interaction of the C4-Fc and E3-Fc constructs with hα7-, hα3β4-, and hα4β2-nAChRs was tested on permeabilized transiently transfected cells to assess binding specificity. To visually pinpoint transfected cells, through fluorescence, either cytoplasmically fused eGFP (hα4-nAChR) or an internal ribosome entry site (IRES) linked eGFP (hα7- and hα3-nAChR subunits) were used (Figure 3c). Both C4-Fc and E3-Fc label cells transiently transfected with hα7-nAChR, while no labeling is observed for hα3β4- and hα4β2-nAChR transfected cells, demonstrating a strong selectivity of both constructs.

Interestingly, sequence analysis determined that both E3 and C4 diverge from the classical nanobody framework in that C4 contains only a single cysteine and thus no disulfide bridge, whereas E3 contains four cysteines, forming the canonical bridge between framework region (FR) 1 and FR3 domains, but also an unusual bridge flanking both extremities of the complementarity-determining region (CDR) 3 (Figure 1a).

### 3/E3 and C4 bind outside of the orthosteric site

To elucidate their binding mode, a competition assay with I^125^ labeled α-Btx, a competitive and highly specific orthosteric antagonist, was performed. HEK293 cells, transfected with the hα7-nAChR/m5-HT_3_A chimeric construct used in panning experiments, were suspended and incubated with 0.01 mg/mL VHH (∼633 nM, with a C-terminal Myc and His-tag) for 2 hours at 4°C and compared to 100 µM ACh (incubated for 1 hour at 4°C) as a control (concentrations are indicated as final concentrations, and all results were normalized to a buffer incubated sample). Scintillation reading of filtered samples, after the addition of a saturating, final, concentration of 5nM of I^125^-α-Btx was added to this mixture, showed that there is no competition between α-Btx and either VHH (Figure 4). This indicates that neither VHH binds to the orthosteric binding site, and therefore bind to an allosteric site elsewhere within the ECD.

**Figure 4:**
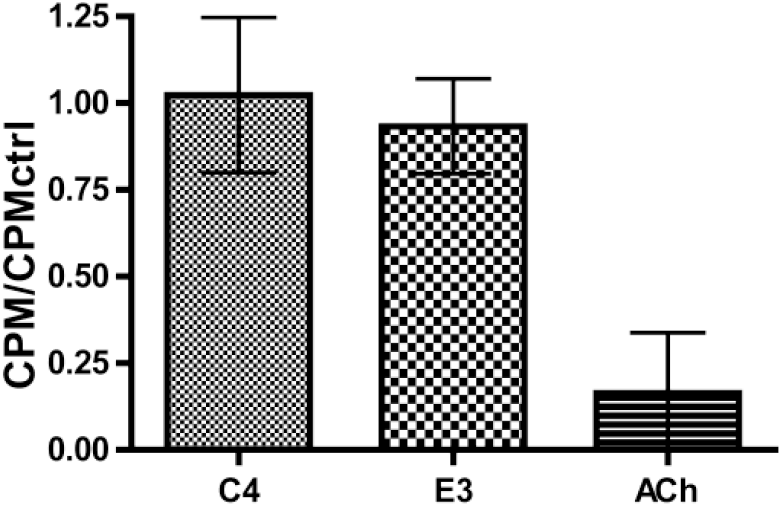
I^125^-αBtx Competition Binding to hα7/m5-HT_3_A. Scintillation counts per minute (CPM) of 5nM I^125^-αBtx bound to hα7-nAChR/m5-HT_3_A transiently transfected HEK293 cells in competition with 100M µM ACh or 0.01mg/mL VHH. Mean of n=6, with error bars showing the standard deviation of the mean.

### 4/E3 is a potent and slowly associating PAM of the α7-nAChR

The functional effect of E3 and C4 (with a C-terminal Myc and His-tag) was investigated by two-electrode voltage clamp electrophysiology on *Xenopus* oocytes expressing the hα7-nAChR (Figure 5). Perfusion of 1 mM ACh, in this methodology, evokes robust currents characterized by a fast onset of activation, a culmination at a peak, and a slower decay of activation whereupon receptor desensitization through prolonged agonist (ACh) application overcomes newly activating receptors, until the agonist is rinsed, and the receptor returns to its resting state. The effects of the nanobodies were investigated at a concentration of 30 µM ACh, corresponding approximately to an EC_10_ concentration to ensure maximal sensitivity of the system.

**Figure 5:**
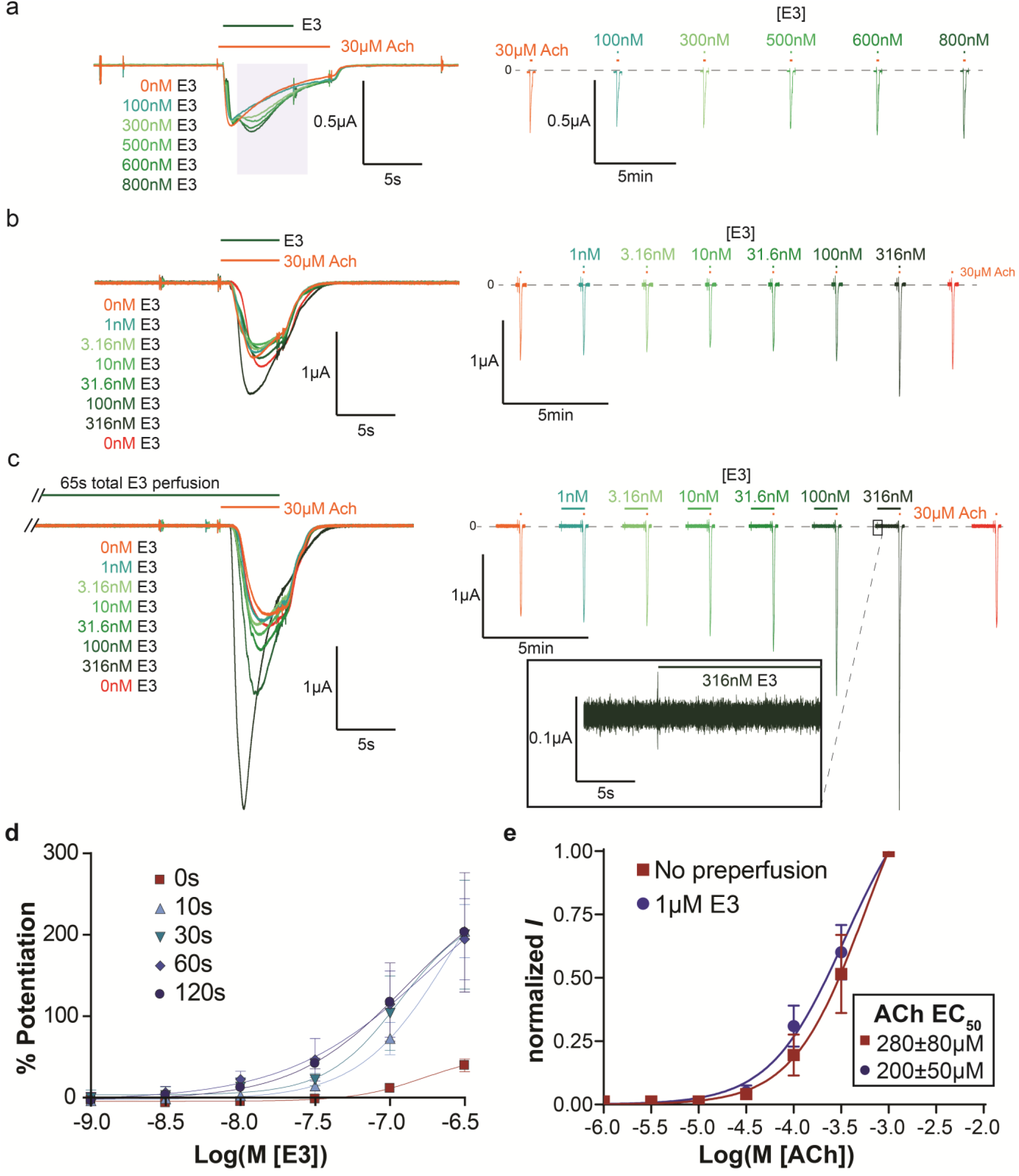
E3-VHH Characterization by Electrophysiology. Representative concentration dependent traces with respective color-coded concentrations of VHH E3 by oocytes injected with a hα7-nAChR-IRES-NACHO plasmid, which allows for the co-expression of hα7-nAChR and NACHO. Superimposed traces (truncated in **c** to allow proper trace visualization), left, are all on the same oocyte and applied chronologically, in the same order as listed. To distinguish the traces the same continuous train of perfusion is shown to the right with an identical current, vertical, scale bar. **a** “Post-perfusion” protocol with E3 co-applied in the middle of a 30µM ACh response. (representative of n=5). Shaded area on the left represents the effective time the receptor is in presence of E3. 30µM ACh response before arrival of E3 in each curve shows that E3 is effectively washed before each new application. **b** “Co-perfusion” protocol with E3 co-applied for the same duration as 30µM ACh (representative of n=5). **c** “Pre-perfusion” protocol using 60s pre-perfusion of E3 with the same concentrations listed in **b** (representative of n=5). Inset on the right shows the beginning of the application of E3 alone on the same time scale as the left panel and vertically zoomed 10-fold. **b** & **c**. 30µM ACh response without E3 (0nM) at the beginning and end of the concentration-response curve shows a complete wash of E3 and stable response of the oocyte. Scale bars are the same size for **b** & **c**. **d** Percent potentiation of 30µM ACh peak response, calculated as [(maximal VHH+ACh current minus maximal 30µM ACh alone current) divided by maximal 30µM ACh alone current] times 100, where 100% potentiation is equivalent to a current two times in amplitude of the 30µM ACh alone current, by various pre-perfusion times of VHH E3 in a concentration-dependent manner (n=5 each). Maximum potentiation is achieved by 60s pre-perfusion with 30s producing only a slightly smaller potentiation, therefore it was decided to keep a 30s pre-perfusion for subsequent experiments. **e** ACh concentration response curves using a 30s “pre-perfusion” protocol with and without a fixed concentration (1µM) of E3. Values are normalized to the peak of 1mM ACh and shown as the mean of n=7 with the error bars displaying the standard deviation. Data show that ACh affinity tends to increase in the presence of E3, but the effect is not significant.

E3 does not elicit any current when applied alone (see Figure 5c right inset for “pre-perfusion” applications of E3 alone before ACh currents). In a first series of experiments, E3 was applied during an application of ACh (“post-perfusion condition”, n=5, Figure 5a). At 800 nM, E3 elicits a clear increase of the ACh current, and the rise-time of this potentiation appears relatively slowly in comparison to the rise time of ACh alone. The potentiation by E3 does not overcome receptor desensitization as seen by a time-dependent decrease during VHH application, indicating that the nanobody favors the activation process but does not preclude the downstream process of desensitization. This potentiating effect is concentration-dependent. To further investigate the kinetics by which E3 modulates the function, several concentration-response recordings were performed where E3 is directly co-applied with ACh (“co-perfusion condition”, n=5, Figure 5b), or where it is additionally perfused for periods of 10, 30, 60 (Figure 5c), and 120s before the ACh/E3 co-application (“pre-perfusion conditions”, n=5 each time-point, Figure 5d). Longer pre-perfusion conditions generated significantly larger potentiation, especially at low nanobody concentrations, where 316 nM E3 obtained near maximal potentiation, indicating that the nanobody associates to the receptor with a slow kinetics in the recording conditions. This slow kinetics is probably the main reason for the moderate potentiation observed in “post-perfusion” conditions. Indeed, in these conditions, E3 will slowly bind and potentiate the receptor, a process during which a significant fraction of the activated receptors will desensitize, generating a net reduction of the peak current. Altogether, the data unambiguously show that E3, with low nanomolar apparent affinity and relatively slow binding kinetics, elicits potentiation of ACh-elicited currents and does not preclude desensitization.

In concentration-response curves, PAM binding usually increases the apparent affinity of the agonist [15]. ACh concentration-response curves in the absence or presence of a 1 µM concentration of E3 were therefore measured. E3 tends to cause a small decrease in the apparent EC_50_ of ACh (n=7, Figure 5e), but this effect is not statistically significant due to the variability of the responses between individual oocytes. It is noteworthy that the kinetics of the rise-time and decay-time of the responses were also variable as illustrated by the current traces shown in Figures 5-7. This precludes a kinetic analysis of the activation and desensitization processes. Further studies would thus be required to investigate quantitatively a possible effect of E3 binding on the kinetics of activation and desensitization.

### 5/C4 acts as a SAM that competes with E3 binding

Conversely, C4 elicited no response on its own nor showed any discernable modification of α7-nAChR function when co-applied with ACh (tested at 316nM with 15s perfusion alone or in “post-perfusion” conditions with ACh n=3 each, data not shown). Interestingly, when using a modified 30s “purely pre-perfusion” procedure (see Methods), the ACh-gated current after pre-perfusion of a mixture of E3 (150 nM) with C4 (at either 150nM or 1.5µM) is near identical to that in the absence of nanobodies (5±10% and -9±21%, in the presence of the respective C4 concentrations, of the % potentiation by 150nM E3 alone); whereas E3 (150 nM) pre-perfused alone on the same oocyte just before the application of these mixtures produced robust potentiation of 30µM ACh (75±17% and 71±28%, n= 3 and 5 resp., Figure 6a). These data indicate that C4, which has no discernable functional effect on the α7-nAChR alone, binds to inhibit the potentiating effect of E3. The simplest interpretation is that the C4 and E3 binding sites overlap at least partially, with C4 inhibiting E3 binding in a competitive manner. Whatever the case, the data allow classification of C4 as a silent allosteric modulator (SAM).

**Figure 6:**
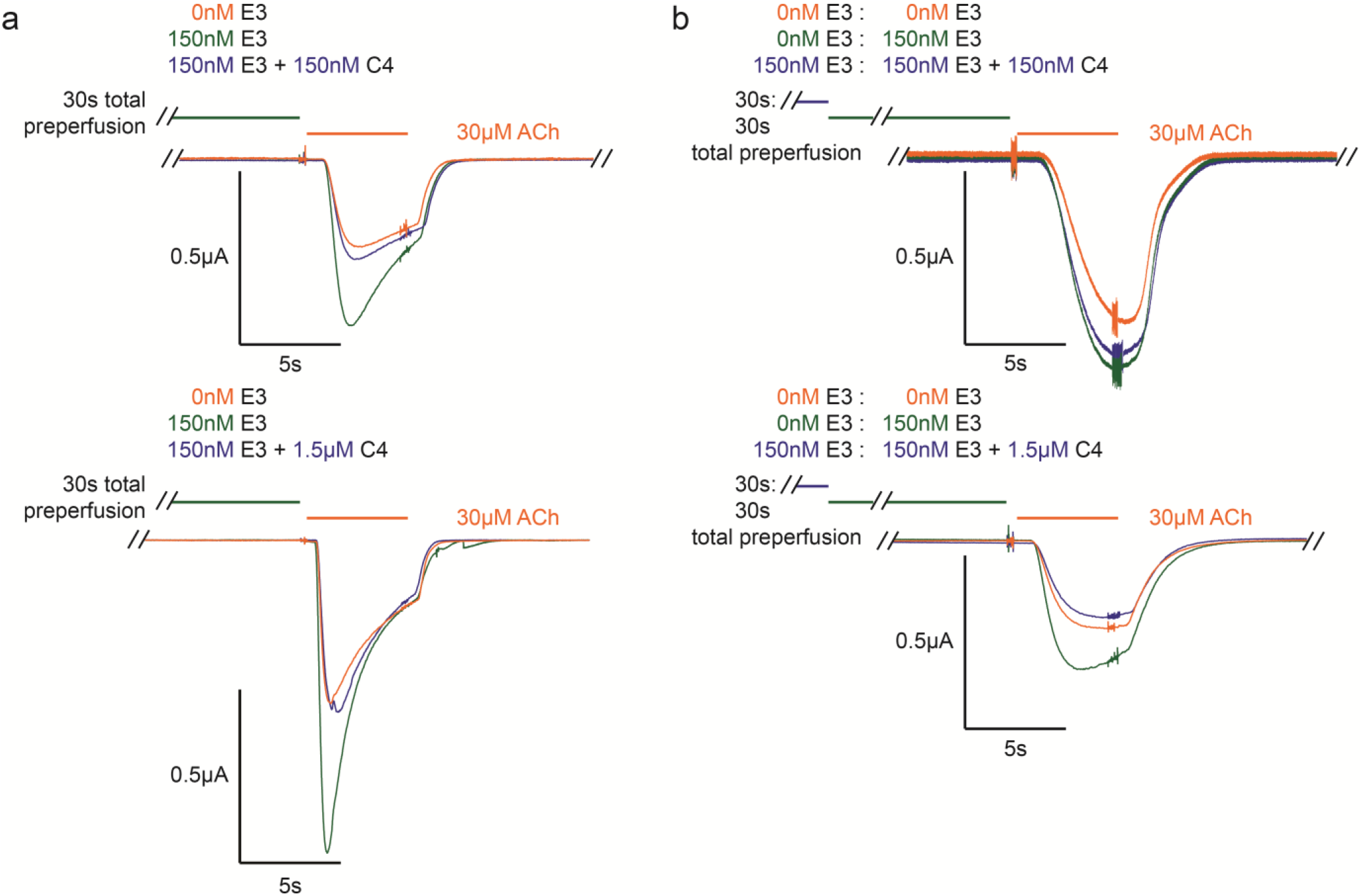
E3-C4 Competition. **a** Top and bottom are representative traces (of n=3 and n=5 resp.) of a mixture of C4 (150nM and 1.5µM, top and bottom resp.) and 150nM E3 using a modified 30s “purely pre-perfusion” protocol, which shows that C4 blocks E3 potentiation. **b** Top and bottom are selected traces (from n=3) of C4 (150nM and 1.5µM, top and bottom resp.) and 150nM E3 competition using a “double purely pre-perfusion” protocol, where 150nM E3 was pre-perfused for 30s before the combined mixture. The concentration-dependent competition strongly suggests that the two VHH’s compete for an overlapping binding site.

To further examine a possible competitive binding mechanism between E3 and C4, a “double purely pre-perfusion” procedure was performed to test whether C4 could displace already bound E3. In this condition, E3 (150nM) is perfused for 30s, then a mixture of E3 (150nM) and C4 (at either 150nM or 1.5µM) is perfused for 30s, followed by activation by 30µM ACh. It should be noted that the potentiation of E3 (150nM), pre-perfused alone for 30s, was less pronounced in these experiments (43±14%, n=6). 1.5µM of C4 was found to out-compete 150nM of pre-bound E3, yielding no potentiation (−14±18% of E3 potentiation, n=3), whereas 150nM C4 only partially blocks the effect of E3 (48±30% of E3 potentiation, n=3) (Figure 6b). The data thus suggest that C4 can displace already bound E3 molecules. Whereas, the reduced ability of C4, at 150nM, to displace already bound E3 is most probably due to the slow binding kinetics of these nanobodies.

It is noteworthy that comparison of the sequence of E3 and C4 show significant conservation at CDR1, CDR2 and half of CDR3, further arguing for overlapping binding sites. Most importantly, these data indicate that this allosteric ECD site is capable of various modulatory actions akin to the orthosteric site.

### 6/Bivalent E3-E3 is a quasi-irreversible PAM

To increase the affinity and avidity of E3 against hα7-nAChR, a bivalent construct was generated, where two E3 nanobodies are separated by a 16-residue flexible linker (Figure 1). E3-E3 displays potentiating properties similar to that of E3 (n=11, Figure 7a & n=4, Figure 7b). Strikingly, the potentiating effect of E3-E3 did not vanish upon prolonged wash out of the oocyte (Figure 7c), achieving 94±12% of the initial maximal potentiating current with repeated applications of 30µM ACh alone out to ∼50 min (for n=4) after the last application of E3-E3. Whereas, the effect of E3 totally vanished upon 3 min of washing (Figure 5). E3-E3 is at least as efficient as E3 to potentiate the ACh-elicited currents, but evaluation of a concentration dependent curve is rendered moot with the impossibility of washing out the construct. Thus, the avidity conferred by the bifunctional character makes E3-E3 a quasi-irreversible potentiator in the oocyte recording system. Like E3, E3-E3 tends to cause a small decrease in the apparent EC_50_ of ACh (n=4, Figure 7b), but this effect is not statistically significant.

**Figure 7:**
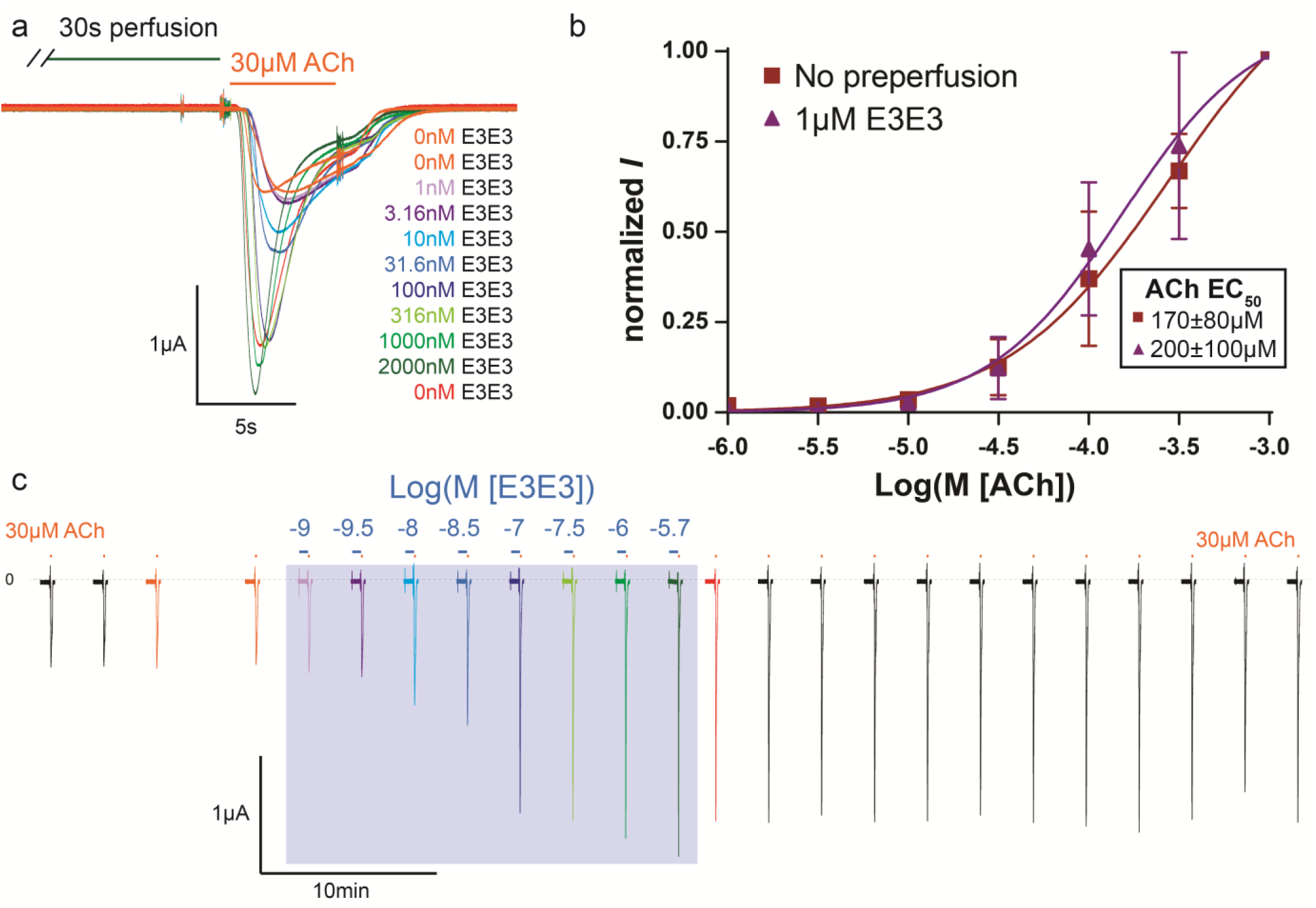
E3-E3 VHH Characterization by Electrophysiology. **a** Representative (of n=11) overlay of concentration dependent traces with respective color-coded concentrations of E3-E3 bivalent VHH using a “purely pre-perfusion” protocol. **b** ACh concentration response curves using a 30s “pre-perfusion” protocol with and without a fixed concentration (1µM) of bivalent E3-E3, where effectively the oocyte is in a constant presence of the bivalent VHH. Values are normalized to the peak of 1mM ACh and shown as the mean of n=4 with the error bars displaying the standard deviation. Data show that ACh affinity tends to increase in the presence of the bivalent E3-E3, but the effect is not significant. **c** An extension of the same traces from A. (similarly color coded) shown in a continuous chronological fashion, where additional applications of 30µM ACh alone are included (not shown in **a**) before and after the concentration-dependent application of E3-E3 bivalent VHH (traces within blue box). Data show that potentiation is maintained even 40 min after last application of the bivalent E3-E3.

## Discussion

The classical technique to generate specific nanobodies against a given target is to immunize alpacas with purified protein, a procedure that requires large quantities of protein (>1mg). Within the pentameric ligand-gated ion channel family, this technique has been successfully applied for members yielding good expression in recombinant systems such as the ELIC [24], the 5-HT_3_ receptor [22], and the GABA_A_ receptor [23]. This article focuses on the α7-nAChR, a subtype that shows low levels of expression in recombinant systems [10]. To avoid the time-consuming overexpression and purification steps, direct immunization with a cell-line transiently transfected with a hα7-nAChR/m5-HT_3_A chimera that has good expression levels was performed. Such a procedure is expected to stimulate the production of a wide range of nanobodies recognizing many proteins present at the surface of the cells. Therefore, a carefully designed panning strategy was completed, yielding, after a few rounds, two nanobodies that bind specifically to the α7-nAChR. The procedure has also the advantage of injecting membrane-inserted protein, ensuring native pentameric assembly of the receptor, increasing the chance to isolate antibodies targeting a properly folded ECD in a pentameric conformation. This procedure should be applicable to other pentameric ligand-gated ion channels with weak expression levels and/or low stability after extraction from the plasma membrane.

The α7-nAChR is activated by choline with an EC_50_ around 500 µM in an oocyte, with choline already producing clearly detectable responses at concentrations of 30 µM or less [28]. Since levels of choline in human plasma have been reported to range between 5 to 15 µM [29], it is possible that such a concentration in the alpaca plasma would stabilize a significant fraction of the receptor on the cells which were injected in the active or desensitized conformational states during immunization. This could direct the generation of a PAM, such as E3, that favors the active state and does not preclude desensitization. The generation of a SAM, in C4, is consistent with this assessment as a VHH that equally binds the resting state and active state would be equally possible in such an environment, as an equal conformational affinity would stabilize neither state, and thereby lack a functional modulation.

Structural and/or mutational studies of the α7-nAChR in complex with the described nanobodies are required to fully understand their mechanism of action. To date, several nanobodies or even fragments of antibodies have been structurally resolved with pentameric ligand-gated ion channels. These structures help gleam possible binding locations for E3 and C4. From these structures, two regions can be directly excluded from the fact that no competition with α-Btx is observed. The first, an exterior site just below the orthosteric pocket, structurally resolved on GABA_A_ [30, 31] and a nAChR [32] and the second, an exterior binding site directly overlapping and slightly above that of α-Btx, found with the 5-HT_3_ receptor [22]. A third site on the complementary subunit to the right of and slightly above the orthosteric pocket, visualized on ELIC [24] and GABA_A_ [33] respectively, is likely occluded in the α7-nAChR due to a unique glycosylation site. This site in the prokaryotic ELIC was found to have PAM effects. It may still be possible that the VHH’s accommodate for a sugar moiety in this region. Finally two apical sites have been observed: the main immunogenic region, bound by an antibody fragment on a nAChR [34], and a single VHH that has multiple interactions with the apex of the ELIC, laying on the top and interacting through the CDRs with multiple subunits in the vestibule [24]. The later was determined to be a NAM and through its binding pose it would be predicted that it would not be conducive to an enhancement of avidity through bivalent binding. Whereas, the former was also found to have a slight increased affinity on agonist binding but no electrophysiological effects at saturating agonist concentrations. Therefore, of the currently resolved binding locations the data would be compatible with two regions, both of which have also hinted to positive modulatory effects. In conclusion, E3 binds outside of and above the orthosteric site, at an apical position. Structural and/or mutational studies are necessary to determine the exact location of VHH binding.

Interestingly, the dimerization of E3 allows the formation of a quasi-irreversible PAM. This striking effect is probably related to an increase of the avidity of this bivalent molecule. In retrospect, this feature probably explains why, in immunofluorescence experiments that involves several rinsing steps after VHH binding, the labeling by monovalent E3 was weak, while that of bivalent E3-Fc was robust. Several examples of dimerization of nanobodies directed against viral proteins have been reported to potentiate the neutralizing activities of the parental nanobody [35–38]. This bivalent construct can serve as a unique therapeutic agent, especially for neurodegenerative disorders where α7-nAChR expression is diminished, potentially permanently amplifying nAChR response signals without having a temporal effect (through modification of binding/desensitization properties) on the nAChR response. This possibility should be further studied to see if adverse effects due to prolonged potentiation of α7-nAChR arise, or if the bivalent nature of the VHH yields problems in administration, such as crossing the blood-brain barrier.

## Conclusions

The two nanobodies C4 and E3 constitute a novel class of allosteric modulators of the α7-nAChR. They show high specificity among nAChR subtypes, a feature characteristic of antibody-antigen recognition that involve a large area of interaction. These nanobodies are easily expressed in milligram amounts in cell-lines and can be easily engineered as illustrated here by the generation of a quasi-irreversible bivalent potentiator. They will be useful for a wide range of applications, notably the investigation of native receptors in brain tissues in immunofluorescence and immunoprecipitation assays. They will also be precious for the investigation of the receptor and as pharmacological tools to help structural studies. Finally, they constitute an original family of allosteric modulators with far reaching potential medical applications such as cognitive enhancers, or as a potential treatment against α7-nAChR auto-antibodies that are found in some patients diagnosed with schizophrenia [39].

## Methods

### Animal immunization

All immunization processes were executed according to the French legislation and in compliance with the European Communities Council Directives (2010/63/UE, French Law 2013-118, February 6, 2013). The Animal Experimentation Ethics Committee of Pasteur Institute (CETEA 89) approved this study (2020-27412). We subcutaneously injected an adult alpaca at days 0, 21, and 28 with approximately 10^8^ hα7-nAChR/m5-HT_3_A chimera-transfected HEK293 cells mixed with Freund complete adjuvant for the first immunization and with Freund incomplete adjuvant for the following immunizations. Repetitive immunogen administrations were applied to stimulate a strong immune response. A blood sample of about 250 mL of the immunized animal was collected and a Phage-Display library in a pHEN6 phagemid vector of about 2×10^8^ different clones was prepared as described before [40], and detailed below.

### Molecular biology

The plasmids for all nAChR constructs were human (h). The hα7-nAChR/m5-HT_3_A plasmid was created merging the human α7-nAChR sequence with the mouse 5-HT_3_A sequence at V202 of the α7-nAChR mature sequence (as described in [26]). pcDNA3.1+ vector was used for all constructs containing an internal ribosome entry site (IRES), which allows for expression of the sequence following the IRES. The expression of this sequence is slightly reduced as compared to the sequence directly behind the promoter sequence and before the IRES. The pMT3 vector was used for the eGFP alone and the hα4-nAChR which contained an ICD linked GFP (hα4_GFP_); where an eGFP sequence was inserted in the cytoplasmic domain between SCK_395_/S_396_PS following the mouse α4-nAChR construct created by Nashmi, R. et al. [41]. The pCMV6-XL5 vector was used for the NACHO construct [42]. Hα7- and hα3-nAChR-IRES-eGFP constructs, used for immunofluorescence experiments, contained the respective nAChR sequence, followed by an IRES element from the encephalomyocarditis virus, and then the coding sequence of eGFP (sequences extracted from pIRES2-EGFP, Clontech). Hβ4_StrepTagII_-nAChR-IRES-mTurquoise, contained the human β4 sequence with a C-terminal StrepTagII the aforementioned IRES element followed by an mTurquoise GFP. For transduction into the CHO-K1 cell-line hα7-nAChR-IRES-eGFP, along with the chaperone protein constructs: NACHO-IRES-mKate and Ric3-IRES-TdTomato, were cloned into a modified pTRIPΔU3 vector with a CMV promoter, and lentiviral particles created as described in the supplementary methods of Maskos, U. et al.[43]. In the case of the hα7-nAChR plasmid used in two-electrode voltage clamp experiments the IRES was followed by the coding sequence for NACHO. Synthetic genes of the anti-α7 nanobody derivatives were designed and purchased from EurofinsGenomics (Ebersberg, Germany), digested using SfiI and NotI and ligated into the pHEN6 phagemid [44].

### -Library creation through PCR amplification of VHH DNA from serum

About 250 mL bloodletting samples of the immunized alpaca in EDTA-coated tubes were collected and inverted twice to inhibit coagulation. Histopaque-1077 (Roche) was employed to separate lymphocytes according to the manufacturer’s instructions. Total RNA was extracted, following the RNeasy minikit (Qiagen) protocol, from isolated lymphocytes and its purity verified using the Agilent RNA 6000 Nano Assay system.

This total RNA was subsequently reverse-transcripted to generate a diverse cDNA library. This cDNA was then amplified by overlap extension PCR which allows for the isolation of the ∼400bp VHH domain. These PCR fragments were also digested using SfiI and NotI and ligated into the pHEN6 phagemid, which was transformed through electroporation into *E. coli* TG1 cells.

### Cell culture

HEK293 and Vero cells were cultured in high glucose Dulbecco’s Modified Eagle Medium supplemented with 110 mg/L sodium pyruvate and 862 mg/L of L-alanyl-glutamine as well as 10% fetal bovine serum (DMEM-FBS) with or without the addition of an antibiotic mixture of penicillin/streptomycin (10 U/ml and 10 μg/ml resp.) at 37°C with 5% CO_2_. Expi293F cells were maintained at 99% viability per manufacturer’s recommendations in Expi293™ Expression Medium (ThermoFisher) at 37°C with 8% CO_2_ under constant agitation and passed every three days. CHO-K1cells were cultured in DMEM/F12 Medium (Gibco) supplemented with 10% FBS and penicillin/streptomycin (10 U/ml and 10 µg/ml) at 37°C with 5% CO2.

### -Generation of the hα7-nAChR-CHO-K1 stable cell line

Lentiviral particles were directly added to CHO-K1 cells during their seeding at 1×10^5^ cells/well respecting a particle ratio of 5:1:1 (hα7-nAChR-IRES-eGFP, NACHO-IRES-mKate, Ric3-IRES-TdTomato resp.) which was achieved using 1µg of p24 for hα7-nAChR particles and 0.2µg for the chaperones. After 3 days of expression, cells were sorted by FACS (FACS-Aria III; BD Life Sciences) and cells expressing the 3 reporter genes were seeded in 96 wells plate to obtain individual clones. Clones expressing a high level of hα7-nAChR at the plasma membrane were selected by α-Btx immunostaining.

### Biopanning by Phage-Display on cells

To produce nanobody-phages, 100 µl of the *E. coli* TG1 library was inoculated into 100 ml of 2xYT medium (Tryptone 16g/L, Yeast Extract 10g/L, NaCl 5g/L, pH 7) supplemented with ampicillin (100 µg/ml) and glucose (1%). The culture was grown at 37°C under shaking at 200 rpm until OD_600_ (optical density at λ600) reached ∼0.5-0.6. Recombinant nanobody-displaying phages were rescued with 2×10^11^ PFU of M13KO7 Helper phage (New England Biolabs). Rescued phages were suspended in 1 ml of phosphate buffered saline (PBS, Gibco) supplemented with bovine serum albumin (BSA, Sigma-Aldrich) at a final concentration of 1% (PBS-BSA).

In a round of panning, 0.5 mL of produced nanobody-phages was added to 4.5 mL of PBS-BSA solution as a phage panning medium. 1×T75 confluent HEK cells (∼10^7^) were washed with PBS and resuspended to individual cells. The suspension was centrifuged at 2000 rpm at 4°C for 15 minutes. Cells were resuspended with 5mL phage panning medium for the first-depletion round, and maintained at 4°C for 1 h with gentle shaking. The suspension was then centrifuged (same conditions as above) where the supernatant phages were recovered. This depletion step was repeated and then completed once more on Vero cells (5 mL, ∼2^7^) to eliminate non-specific nanobodies directed against common membrane proteins. Phages were then finally incubated with hα7-nAChR/m5-HT_3_A chimera-transfected Vero cells, again in 5 mL at 4°C for 1 h with gentle shaking to select target-specific nanobody-phages. After incubation, suspension was centrifuged (as above) and washed 10 times with PBST (PBS with 0.01% Tween 20) (each time for 10 minutes), followed by 2 washes with PBS to remove Tween 20. The cells were then lysed with 1 mL of 100 mM triethylamine (Sigma-Aldrich) for 5 minutes at 4°C and pH was neutralized with 0.5 mL of 1M Tris (pH 7.4). Eluted phages (in triethylamine lysate) were amplified by infecting exponentially growing *E. coli* TG1 at OD_600_ of ∼0.5 to propagate phages for the next round of selection.

### Enzyme-linked immunosorbent assay (ELISA)

96 isolated clones from the second and the third rounds of panning were tested for nanobody-phage specific binding to α7-nAChR in cell-ELISA.

hα7-nAChR/m5-HT_3_A chimera-transfected Vero cells (10^5^ cells/mL) were coated onto 96-well plates pretreated with poly-D-Lysine (Sigma-Aldrich), according to manufacturer’s recommendations. A negative control (non-transfected Vero cells) was prepared under the same conditions. Cells were then fixed by addition of 4% (w/v) paraformaldehyde (Sigma-Aldrich) in PBS for 20 min, culture were then rinsed three times with PBS and treated with 3% (w/v) hydrogen peroxide (BioRad) in PBS for 5 min to minimize endogenous peroxidase activity, and finally washed twice in PBS. Plates were then blocked with PBS-BSA for 1h at 37°C.

Nanobody-phage unique clones were separately grown in 300 µl of 2xYT medium supplemented with ampicillin (100 µg/ml) in a 96-deep wells plate, unique nanobody-phage’s production was induced by co-infection with the Helper Phage and culture was left overnight at 30°C under shaking. The supernatant was retrieved by centrifugation (2500 rpm/20 minutes at 4°C) and the produced unique phages were tested in parallel for their specific binding on hα7-nAChR/m5-HT_3_A chimera-transfected Vero cells and on non-transfected ones as a control. The supernatant was diluted to 1/5 in PBS-BSA, and 100 µl of the dilutions were applied to fixed cells in corresponding wells, after incubation for 1h at 37°C, plates were washed 3 times with PBST, and bound nanobody-phages were labelled by the addition of anti-M13 antibody [HRP] (SinoBiologicals). After 5 washes with PBST, HRP substrate reagent (OrthoPhenyleneDiamine, Sigma-Aldrich), prepared according to manufacturer’s recommendation, was added. The reaction was stopped by the addition of 50 µl per well of 3M HCl, and the absorbance of the samples was measured at 492 λ in a spectrophotometer-microplate reader (Sunrise, Tecan). Samples with an absorbance ratio (positive/control) exceeding 3 were considered as α7-positive clones. These were then sequenced, and the sequences were analyzed.

### Anti-α7-nAChR nanobody engineering, expression, and labeling

#### -Engineering

The pHEN6 vector used for library construction allows for direct bacterial periplasmic expression and purification of selected nanobodies with a C-Myc tag and a 6xHis tag at their C-terminal (VHH in Figure 1).

The gene coding for a monovalent nanobody with a 6xHis tag in the N-terminal and an extra C-terminal Cys-Ser-Ala motif (CSA) enables site-specific labeling of the nanobody via maleimide (VHH-CSA in Figure 1) [45]. The gene for the bivalent derivative is composed of two molecules of the same nanobody linked together by a flexible (GGGGS)_3_ linker, with a C-Myc tag and a 6xHis tag at the C-terminal (VHH-VHH in Figure 1). The VHH-Fc gene had the C-terminal linker listed in Figure 1.

#### -Expression

Genes were cloned into the pHEN6 vector for periplasmic expression in the *E. coli* strain: BL21(DE3). Proteins were produced as periplasmic components in 1 L of the NZY Auto-Induction TB medium (NZytech) according to the manufacturer’s recommendations and were purified by immobilized metal affinity chromatography on a HiTrap TALON® crude 1 mL column (Cytiva). After extensive washings with PBS containing 150 mM NaCl (PBS/NaCl), proteins were eluted in PBS/NaCl buffer supplemented with 500 mM imidazole. Bacterial production yields varied from 1-12 mg/L of culture.

Alternatively, the nanobodies’ engineered genes were cloned into a pFUSE-derived vector (InvivoGen); this vector harbors a human IgG1-Fc domain. Consequently, the nanobody was expressed as a Fc-fusion bivalent antibody. The vector was used to transfect Expi293F mammalian cells (ThermoFisher), and protein expression was carried out according to manufacturer’s recommendations. Protein was then purified from the expression medium by affinity chromatography on a 1 mL protein G column (Cytiva). After sample application, the column was washed with 20 column volumes of PBS and the protein was subsequently eluted with 10 column volumes of PBS supplemented with 0.1 M Glycine (pH=2.3). Production yields were above 25 mg/L of culture.

#### -Labeling

Site specific labeling of the engineered nanobody was done using maleimide Alexa Fluor™ 647 C2 or Alexa Fluor™ 488 (ThermoFisher). Briefly, the anti-α7 nanobody with additional CSA motif was surrendered to a mild reduction by adding 10 molar excesses of tris(2-carboxyethyl) phosphine (TCEP) (Sigma-Aldrich) at 25°C for 30 minutes. Reduced protein was then incubated for 2 hours with 20 molar excess of the respective maleimide Alexa Fluor dissolved in DMSO. Labeled protein was filter-dialyzed against 10 L of PBS using a 3K-CutOff Slide-A-Lyzer™ Dialysis Cassette (ThermoFisher). Labeling quality was assessed by Mass Spectrometry as previously described [46].

### Flow-Cytometry

1-5×10^6^ hα7-nAChR/m5-HT_3_A transfected cells, with >90% viability, were centrifuged at 1000 *g* for 10 min at 4°C, and resuspended to obtain a final concentration of 100-300cells/µL in ice cold PBS solution supplemented with 10% FBS and 1% NaN_3_ to prevent the alteration and internalization of surface antigens, which can induce a loss of fluorescence intensity. 100 μL of this cell suspension was added to each well in a 96-well U-bottom plate, and incubated for 30min with the addition of 0.02mg/mL or 0.1mg/mL of a tested VHH (final concentrations) at room temperature or PBS for positive control. Cells were centrifuged at 1000 g for 5 min at 4°C, the supernatant was removed and 200 μL ice-cold PBS added. This wash was repeated three times. Cells were then incubated for 30min, covered to keep plates in the dark, at room temperature with 1μg/mL of anti-C-Myc tag AlexaFluor488 conjugated antibodies (Abcam), or 0.1mg/mL of AlexaFluor488 conjugated α-Btx (Abcam) for the positive control (final concentrations). Cells were centrifuged at 1000 g for 5 min at 4°C, the supernatant was removed and 200 μL ice-cold PBS added. This wash was repeated three times, with the final resuspension in ice-cold PBS, 1% NaN_3_ with 3% BSA. Flow cytometry analyses were performed on a cytoflex S (Beckman Coulter) with the Cytexpert software version 2.2. FITC was read with the laser wavelength of 488λ, using the filter 525/40, whereas PE with a laser 581λ and the filter 586/42.

### Immunofluorescence

Transduced CHO-K1 cells expressing hα7-nAChR and chaperone proteins were cultured on 50µg/ml poly-D-lysine (Sigma-Aldrich) and 330µg/ml Collagen type 1 (Sigma-Aldrich) coated glass coverslips. 48h later, the cells were fixed with 2% paraformaldehyde (PFA) for 15 min followed by blocking of non-specific binding, with 3% BSA in PBS for 1h. The nanobodies were diluted to 5µg/ml with PBS-BSA and incubated with the coverslips for 2h at room temperature.

HEK293 cells were cultured on poly-D-lysine (Sigma-Aldrich) coated glass coverslips, according to manufacturer’s recommendations. These cells were transfected using 10 μg DNA and the JetPrime transfection reagent (Polyplus), again according to manufacturer instructions. 48h-36h after transfection, cells were fixed with 4% PFA and for permeabilized cells incubated for 2 min with a solution of ethanol+methanol (1:1). Non-specific binding was blocked with 10% BSA in PBS for 5 min at room temperature. The nanobodies were diluted to 5µg/ml also with 10% BSA in PBS and incubated with the coverslips for 2h at room temperature.

The hα7-IRES-eGFP and NACHO plasmids were transfected together (2:1 ratio), the hα3-IRES-eGFP and hβ4_StrepTagII_-IRES-mTurquoise were transfected together (1:1 ratio), and a plasmid containing the hβ2-nAChR subunit was transfected together with the hα4_GFP_ construct (1:1 ratio). Proper expression of the respective nAChRs was verified by an anti-StrepTagII antibody for the hα3β4-_StrepTagII_-nAChR, by the fused GFP for the hα4_GFP_β2-nAChR, and with an Alexa647 labeled α-Btx for the hα7-nAChR (Supp Figure 1). Anti-StrepTagII antibody and α-bungarotoxin-Alexa647 (ThermoFisher) were diluted in PBS-BSA.

Coverslips were mounted on slides after Prolong-DAPI staining (Invitrogen) and visualized using epi-fluorescence at constant exposure times. Anti-human IgG and anti-mouse IgG coupled to Alexa647 (ThermoFisher) were diluted in PBS-BSA. All experiments were reproduced ≥4 times.

### Radioactive I^125^-α-Btx competition assay

HEK-293 cells were seeded, 48 h before transfection, at a density of 5×10^5^ per 100mm dish in DMEM-FBS supplemented with penicillin/streptomycin (10 U/ml and 10 μg/ml resp.). When cells were approximately 70%-80% confluent, they were transfected with 10µg of the hα7-nAChR/m5-HT_3_A chimeric construct using the JetPrime reagent (Polypus) according to the manufacturer’s protocol, and incubated for another 48 h. Cells were washed and detached using a cell dissociation solution non-enzymatic (non-trypsin) (Sigma-Aldrich). Cells were resuspended to individual cells in 2mL of binding buffer (10mM HEPES, 2.5mM CaCl_2_, 2.5mM MgCl_2_, 82.5mM NaCl, pH 7.2) in the presence of cocktail protease inhibitor (one tablet of cOmplete per 50 mL [Sigma-Aldrich]). Three replicates of 150µL from this resuspension were pipetted to glass vials for each sample. 25µL of each tested VHH (0.1mg/mL) was added with 25µL binding buffer in the control and ACh samples. After an hour incubation at 4°C under agitation, 25 µL of binding buffer was added to all samples and 25 µL of 1mM ACh was added to the ACh sample which were all incubated under agitation for 1h at 4°C. Finally, 50 µL of 25 nM I^125^-Tyr54-α-Btx was added, the samples were well mixed and incubated for 2 h at 4°C. This resulted in final concentrations of 556 nM E3 VHH, 530 nM C4 VHH, or 100µM ACh, and 5nM I^125^-α-Btx. Glass microfiber papers, blocked using 5% skim milk for at least 30min, were then used to vacuum-filter the samples and counts per minute measured using a Berthold LB2111 machine.

### Two-electrode voltage-clamp electrophysiology

*Xenopus laevis* oocytes (EcoCyte Bioscience, Germany and Centre de Resources Biologiques– Rennes, France) kept in Barth solution buffer (87.34 mM NaCl, 1 mM KCl, 0.66mM NaNO_3_, 0.75 mM CaCl_2_, 0.82mM MgSO_4_, 2.4 mM NaHCO_3_, 10 mM HEPES pH 7.6) were nucleus injected with ∼5-8 ng of hα7-IRES-NACHO and∼1ng eGFP plasmids and stored at 16°C-18°C for 48-96 h.

Current recordings were performed using: Axon Digidata 1550A digitizer (Molecular Devices), Axon Instruments GeneClamp 500 amplifier (Molecular Devices), a self-built automated voltage controlled perfusion system, which controls an 8-port and a 12-port electric rotary valve (Bio-Chem Fluidics) both connected to a 2-way 4-port electric rotary valve (Bio-Chem Fluidics) which was flush with the recording chamber entrance allowing for rapid solution application and clean solution exchange, and finally pClamp 10.6 software (Molecular Devices). Electrode pipettes were pulled using a Narishige PC-10 puller. GFP positive oocytes were voltage-clamped at -60mV, unless otherwise noted, and perfused by Ringer’s solution buffer (100 mM NaCl, 2.5 mM KCl, 10 mM HEPES, 2 mM CaCl_2_, 1 mM MgCl_2_, pH 7.3), with sampling at 5kHz, unless otherwise noted. ACh and size-exclusion purified VHH’s were diluted in Ringer’s buffer as specified. ClampFit 10.6 (Molecular Devices) was used for trace analysis and GraphPad Prism 4 (GraphPad Software) for data plotting and statistical analysis. Data are shown as mean ± standard deviation, where each n represents a different oocyte.

The mechanical turning of the valves in the perfusion system produces a noise seen during the recordings, which was not filtered out, allowing for easy recognition of solution exchange times. The main valve allows for one solution to flow to the recording chamber and at the same time the other connected valve, if open, flows to waste. The main 2way/4-port valve exchange time was about 200ms and there is around a 500ms delay between the end of the exchange and arrival of the solution to the oocyte. Solution to be perfused to the chamber was started at least 4s before in the direction of waste to ensure proper rinsing of valve tubing (unless noted otherwise).

#### *“Pre-perfusion”* protocol

The flow of the denoted concentration of VHH applied 6s after the start of the sweep to the recording chamber allowing for a proper analysis of the leak/background current. 10-120s later the main valve was switched to flow a solution that contained the same concentration of VHH with 30µM ACh (or the denoted concentration in the case of ACh concentration-response curve). The valve flowing the VHH alone solution (now to waste) was switched to Ringer’s solution shortly after. Five seconds later the main valve was changed back to flow the Ringer’s solution to the chamber and the other side valve was closed with the sweep finishing recording eight seconds later. There was a three-minute start to start interval between all sweeps which continued washing with Ringer’s solution.

#### *“Purely pre-perfusion”* protocol

Once it was determined that the VHH’s had a slow on and offset time in order to conserve product and simplify solutions, the pre-perfusion protocol was changed to have only a 30µM ACh solution applied for the five second activation after pre-perfusion of VHH in Ringer’s solution, rather than the mixture of ACh and VHH described above.

#### *Modified “purely pre-perfusion”* protocol

Oocytes were voltage clamped at -70mV (for 1.5µM C4) and -80mV (for 150nM C4) and sampled at 2kHz. The flow was switched, without pre-rinsing, to apply a mixture of VHHs at the indicated concentrations directly at the beginning of the sweep for 30s, directly, without pre-rinsing, switching to a solution with 30µM ACh alone applied for five seconds and finishing the sweep with an 8.5s wash (for 1.5µM C4) and a 60s wash (for 150nM C4). The next sweep began directly after the termination of the current sweep.

#### *“Double purely pre-perfusion”* protocol

Oocytes were voltage clamped at -80mV and sampled at 2kHz. Valve switching in all cases was done without pre-rinsing times. For the first sweep 60s of wash buffer was perfused before switching to a solution with 30µM ACh alone for 5s, the subsequent sweep had 30s of wash buffer then switching to apply VHH E3 at 150nM for 30s, before the 30µM ACh alone application, and finally the last sweep had E3 at 150nM directly at the beginning of the sweep, then switched to a solution mixture containing E3 at 150nM along with indicated concentrations of C4 for another 30s, followed by 30µM ACh alone. All sweeps finished with a 56s wash, and the next sweep began directly after the termination of the current sweep.

#### *“Co-perfusion”* protocol

The flow of denoted concentrations of VHH mixed with 30µM ACh was applied 10s after the start of the sweep for a duration of 5s, followed by 15s of recorded wash, with a three-minute start to start sweep interval.

#### *“Post-perfusion”* protocol

In order to properly switch solutions just after the on-set of desensitization there was not enough time to rinse solutions and therefore the main valve could not be used for switching. As a result, the solution was switched between 30µM ACh to 30µM ACh with the denoted concentration of VHH in the secondary valve where the delay of solution arrival is around 1.2s. Therefore, the main valve was switched to flow 30µM ACh 10s after the start of the sweep, with the secondary valve switching to a solution of 30µM ACh with VHH 0.5s later. This achieved the arrival of the VHH around 2s (including valve exchange time) after application of ACh alone. The combination was perfused for 6s before switching back to ACh alone, and the main valve was switched back to Ringer’s after 3s effectively creating a 10s application of 30µM ACh which included a combination of ACh with VHH for 6s directly in the middle. There was a 5min 20s start to start sweep interval which washed the oocyte.

## Acknowledgements

We thank Sylvie Bay for help in nanobody characterization by mass spectroscopy, and Pierre-Henri Commere for help in flow-cytometry experiments. The work and QL were supported by a grant from LanZhou Institute of Biological Products Co., Ltd, LanZhou, China (Institut Pasteur PPU program and doctoral school ED3C). The work was also supported by the Institut Pasteur “Programme transversal de recherche” (PTR 03-17 Nicobinder). ÁN received funding from the “Agence Nationale de la Recherche” (Grant ANR-17-CE11-0030, Nicofive), GDB received funding from the “Institut National du Cancer” INCa, MP and PJC received funding from the European Research Council (Grant #: 788974, Dynacotine), and finally SP and UM were supported by “Fondation pour la Recherche Médicale” FRM-Equipe, INCa, ERA-NET Neuron, and “Centre National de la Recherche Scientifique”.

## List of Abbreviations

5-HT_3_A: 5-hydroxy tryptamine type 3 subunit A
α-Btx: α-Bungarotoxin
ACh: acetylcholine
Ago-PAM: positive allosteric modulator with agonist properties
BSA: bovine serum albumin
cDNA: complementary deoxyribonucleic acid
CDR: complementarity-determining region
cryo-EM: cryogenic electron microscopy
DMEM: Dulbecco’s modified Eagle medium
DMSO: dimethyl-sulfoxide
ECD: extracellular domain
EDTA: ethylenediaminetetraacetic acid
ELIC: *Erwinia* ligand-gated ion channel
ELISA: enzyme-linked immunosorbent assay
FBS: fetal bovine serum
GABA: γ-aminobutyric acid
GFP: green fluorescent protein
hα7-nAChR/m5-HT_3_A: human α7-nAChR/mouse 5-HT_3_A
HEK: Human embryonic kidney
HRP: horse radish peroxidase
ICD: intracellular domain
Ig: immunoglobulin
IRES: internal ribosome entry site
mRNA: messenger ribonucleic acid
nAChR: nicotinic acetylcholine receptor
NAM: negative allosteric modulator
OD: optical density
PAM: positive allosteric modulator
PBS: phosphate buffered saline
PBS-BSA: PBS with 1% BSA
PBST: PBS with Tween 20
PCR: polymerase chain reaction
PFA: paraformaldehyde
resp.: respectively
SAM: silent allosteric modulator
TCEP: tris(2-carboxyethyl) phosphine
TMD: transmembrane domain
VHH: variable domain on heavy chain.

## Authors’ contributions

QL: helped design and performed and analyzed: biopanning, ELISA, VHH (excluding VHH-Fc constructs) engineering, expression, labeling, flow-cytometry, immunofluorescence, radioactivity, electrophysiology experiments; and helped edit the manuscript.

ÁN: designed electrophysiological experiments, designed and constructed recording chamber and perfusion system, analyzed all data, wrote and edited the manuscript.

GA: designed and analyzed biopanning, ELISA, VHH expression and labeling and engineered VHHs; performed alpaca immunization and VHH expression and labeling; and edited manuscript.

GDB: performed electrophysiological experiments involving C4-E3 competition.

MP: performed immunofluorescence experiments and helped edit the manuscript.

SP: designed and performed immunofluorescence experiments and helped edit the manuscript.

NB: helped with radioactivity and immunofluorescence experiments.

RB: performed VHH-Fc expression and helped with immunofluorescence experiments

UM: designed immunofluorescence experiments

PL: performed alpaca immunization and helped design biopanning, ELISA, VHH expression and labeling along with engineering VHHs; helped edit the manuscript.

PJC: conceived project, acquired funding, wrote and edited the manuscript

All authors read and approved the final manuscript

## Data Availability

The vast majority of the data generated or analyzed during this study are included in this published article and its supplementary information file. Any remaining data for the current study is available from the corresponding author(s) on reasonable request.

## Ethics approval and consent to participate

Not applicable

## Consent to participate and Consent to publish

Not applicable

---

**Supplementary Figure 1.**
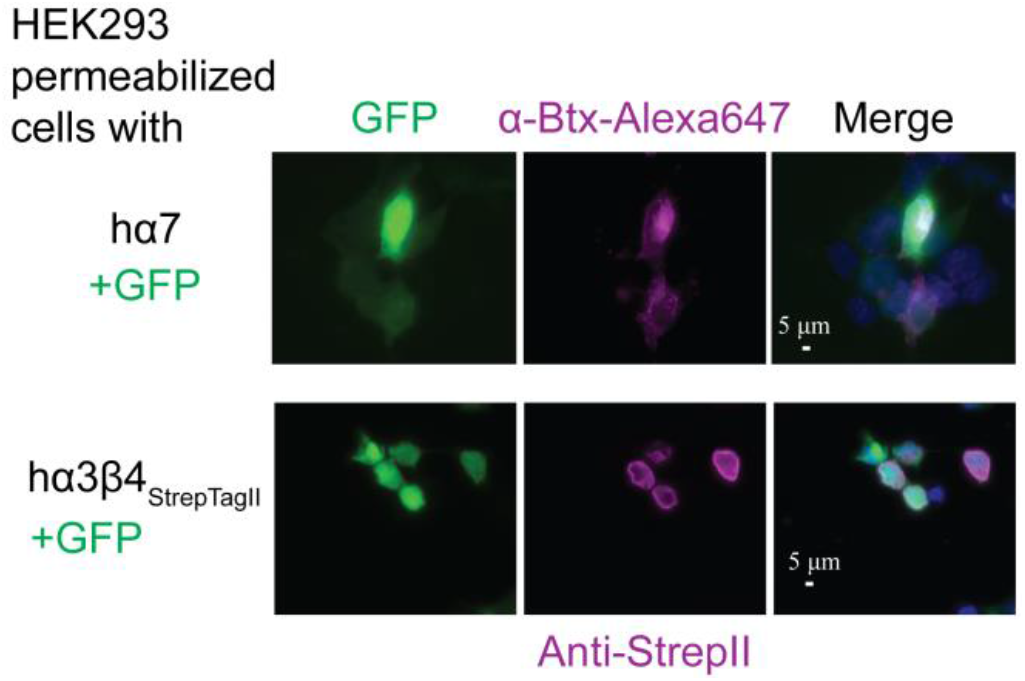
Immunofluorescence expression controls Representative (of n=4) images of permeabilized HEK 293 cells expressing hα7- (top) and hα3hβ4StrepII- (bottom) nAChRs immunostained using conjugated α-Btx-Alexa647 and an anti-StrepII tag detected by a conjugated anti-mouse IgG-Alexa647 respectively. Dapi, shown in blue, stains the cells’ nucleus; Alex647, a red wavelength, is colored as magenta. Cytoplasmic eGFP indicates efficiently transfected cells. Identical exposure times were used to visualize each channel on all conditions.

